# A novel Rab11-Rab3a cascade required for lysosome exocytosis

**DOI:** 10.1101/2021.03.06.434066

**Authors:** Cristina Escrevente, Liliana Bento-Lopes, José S Ramalho, Duarte C Barral

**Author notes:** Correspondence should be sent to: Duarte C Barral, CEDOC, NOVA Medical School|Faculdade de Ciências Médicas, Universidade NOVA de Lisboa, Campo dos Mártires da Pátria 130, 1169-056, Lisboa, Portugal. These authors contributed equally to this work (Cristina Escrevente, Liliana Bento-Lopes).

## Abstract

Lysosomes are dynamic organelles, capable of undergoing exocytosis. This process is crucial for several cellular functions, namely plasma membrane repair. Nevertheless, the molecular machinery involved in this process is poorly understood.

Here, we identify Rab11a and Rab11b as regulators of calcium-induced lysosome exocytosis. Interestingly, Rab11-positive vesicles transiently interact with lysosomes at the cell periphery, indicating that this interaction is required for the last steps of lysosome exocytosis. Additionally, we found that the silencing of the exocyst subunit Sec15, a Rab11 effector, impairs lysosome exocytosis independently of the exocyst complex, suggesting that Sec15 acts together with Rab11 in the regulation of lysosome exocytosis. Furthermore, we show that Rab11 binds the guanine nucleotide exchange factor for Rab3a (GRAB) and also Rab3a, which we described previously as a regulator of the positioning and exocytosis of lysosomes.

Thus, our studies suggest that Rab11-positive vesicles transport GRAB to activate Rab3a on lysosomes, establishing a Rab11-Rab3 cascade that is essential for lysosome exocytosis.

## Introduction

Lysosomes are membrane-bound organelles that play a key role in the degradation of intracellular material. These organelles have long been regarded as the end-point of the endocytic pathway in eukaryotic cells. However, it is now recognized that there are distinct populations of lysosomes in the cell and that lysosomes in the perinuclear region are more acidic than lysosomes at the cell periphery (Johnson et al., 2016). Moreover, lysosomes that localize near the plasma membrane can undergo regulated exocytosis, similar to lysosome-related organelles (LROs) (Andrews, 2000; Rodríguez et al., 1997). In a first calcium-independent step, lysosomes translocate from the perinuclear region to the cell periphery along microtubules, in a process dependent on the multisubunit complex BORC/Arl8/*Salmonella*-induced filaments (Sif)A and kinesin-interacting protein (SKIP) and kinesin-1 (Pu et al., 2015; Rosa-Ferreira and Munro, 2011). In a second step, the pool of pre-docked lysosomes fuses with the plasma membrane, in response to an increase in intracellular calcium. This fusion requires the calcium-sensor Synaptotagmin VII, as well as the R-soluble N-ethylmaleimide-sensitive factor attachment protein receptor (R-SNARE) Vesicle-associated membrane protein (VAMP)-7 on the lysosome membrane and the Q-SNAREs Synaptosomal-associated protein (SNAP)-23 and Syntaxin-4 on the plasma membrane (Martinez et al., 2000; Rao et al., 2004).

Lysosome exocytosis occurs in many cell types and is involved in a variety of cellular processes, including plasma membrane repair and extracellular matrix remodeling and/or degradation (Appelqvist et al., 2013; Hämälistö and Jäättelä, 2016; Jaiswal et al., 2002; Reddy et al., 2001; Samie and Xu, 2014). Plasma membrane repair is particularly important in tissues susceptible to mechanical or ischemic stress, such as skeletal and cardiac muscle (Cheng et al., 2014; Zhao, 2012). Extracellular matrix degradation is frequently enhanced in carcinomas as a result of abnormal lysosomal secretion by cancer cells and leads to stimulation of angiogenesis, tumor growth and cancer cell invasion (Appelqvist et al., 2013; Kallunki et al., 2013; Machado et al., 2015).

Although the molecular machinery involved in lysosome exocytosis is partially known, the role of Rab small GTPases in this process remains poorly elucidated. Recently, we performed a screen to identify Rab proteins that regulate lysosome exocytosis (Encarnação et al., 2016). Rab3a, which is involved in regulated secretion, was one of the most robust hits. Indeed, we found that Rab3a plays an important role in lysosome positioning and plasma membrane repair, as part of a complex formed by the non-muscle myosin heavy chain IIA (NMIIA) and Synaptotagmin-like protein 4a (Slp-4a). Another hit of the screen was Rab11, a Rab protein known to regulate endocytic recycling traffic (Ullrich et al., 1996; Welz et al., 2014). The Rab11 subfamily is formed by Rab11a and Rab11b, which are ubiquitously expressed and Rab11c/Rab25, which is expressed in epithelial tissues of the gastrointestinal mucosa, kidney and lung (Goldenring et al., 1993; Kelly et al., 2012). Interestingly, previous studies have proposed a link between the endocytic recycling pathway and exocytosis in different cell types, including cytotoxic T-cells and bladder umbrella cells (Khandelwal et al., 2013; Ménager et al., 2007). Furthermore, our group established that Rab11b regulates the exocytosis of melanosomes, which are also LROs (Tarafder et al., 2014). However, despite all the evidence, the role of the endocytic recycling pathway and in particular Rab11 subfamily in lysosome exocytosis remains poorly understood.

Similar to other small GTPases, Rab11 proteins cycle between active GTP-bound and inactive GDP-bound forms. GTP hydrolysis is catalyzed by GTPase-activating proteins (GAPs), whereas the exchange of GDP for GTP is promoted by guanine nucleotide exchange factors (GEFs) (Welz et al., 2014). In its active form, Rab11 recruits effector proteins. Among the known Rab11 effectors described in the literature are the Rab11-family of interacting proteins (FIPs) (Baetz and Goldenring, 2013; Hales et al., 2001; Horgan and McCaffrey, 2009; Junutula et al., 2004), motor proteins such as Myosin Va/b (Lapierre et al., 2001; Lindsay et al., 2013) and Sec15, a subunit of the exocyst complex required for the tethering of vesicles to membranes (Wu et al., 2005; Zhang et al., 2004). Moreover, Rab11 was found to interact with other proteins, including Rabin8 (Bryant et al., 2010) and the GEF for Rab3a (GRAB) (Horgan et al., 2013).

In this study, we explored the role of the endocytic recycling pathway in calcium-triggered lysosome exocytosis. We show that the silencing of the small GTPases Rab11a or Rab11b impairs lysosome exocytosis, upon treatment with the calcium ionophore ionomycin. Rab11-positive vesicles were found to transiently interact with lysosomes at the cell periphery, indicating that this interaction is required for the last steps of lysosome exocytosis. Moreover, the silencing of the exocyst subunit Sec15, a Rab11 effector, impairs lysosome exocytosis, independently of the exocyst complex. Furthermore, Rab11 co-immunoprecipitates with Rab3a and the silencing of GRAB, a Rab11 interacting partner and a Rab3a GEF was also found to impair lysosome exocytosis. Thus, our findings describe for the first time a Rab11-Rab3a cascade required for lysosome exocytosis.

## Results

### Rab11a and Rab11b are required for calcium-dependent lysosome exocytosis

To identify Rab proteins that regulate calcium-dependent lysosome exocytosis, we performed a lentiviral shRNA screen, in THP1 monocytic cells, targeting 58 different Rab GTPases (Encarnação et al., 2016). The screening identified several Rabs potentially involved in lysosome exocytosis, including Rab3a and Rab10 (Encarnação et al., 2016; Vieira, 2016), as well as Rab11a and Rab11b. Rab11 proteins are known to regulate the endocytic recycling pathway. Interestingly, Rab11a and Rab11b have been described to be involved in LRO exocytosis, but the molecular mechanism is poorly understood (Ménager et al., 2007; Tarafder et al., 2014; Van der Sluijs et al., 2013).

To confirm that Rab11a and Rab11b play a role in calcium-dependent lysosome exocytosis, we silenced Rab11a or Rab11b isoforms, in HeLa cells and lysosome exocytosis was stimulated with the calcium ionophore ionomycin. Upon lysosome exocytosis, the lysosomal membrane fuses with the plasma membrane. Therefore, the expression of the late endosome/lysosomal marker lysosome-associated membrane protein 1 (LAMP1) on the cell surface was measured by flow cytometry, using an antibody that specifically recognizes the luminal domain of the protein. Non-viable cells were excluded by staining with propidium iodide (PI). We observed that HeLa cells silenced for Rab11a or Rab11b using different shRNA hairpins for each, show impaired cell surface expression of LAMP1, upon ionomycin stimulation, when compared with cells transduced with lentiviruses containing an empty vector (Empty) or encoding a non-targeting shRNA (Mission) (Fig. 1A). From the five hairpins tested (Table S1), four that effectively silenced the protein, gave similar results and therefore only two representative ones are shown. Quantitative RT-PCR and Western-blot confirmed that Rab11a and Rab11b were specifically and efficiently silenced (Fig. S1A-D).

**Fig. 1.**
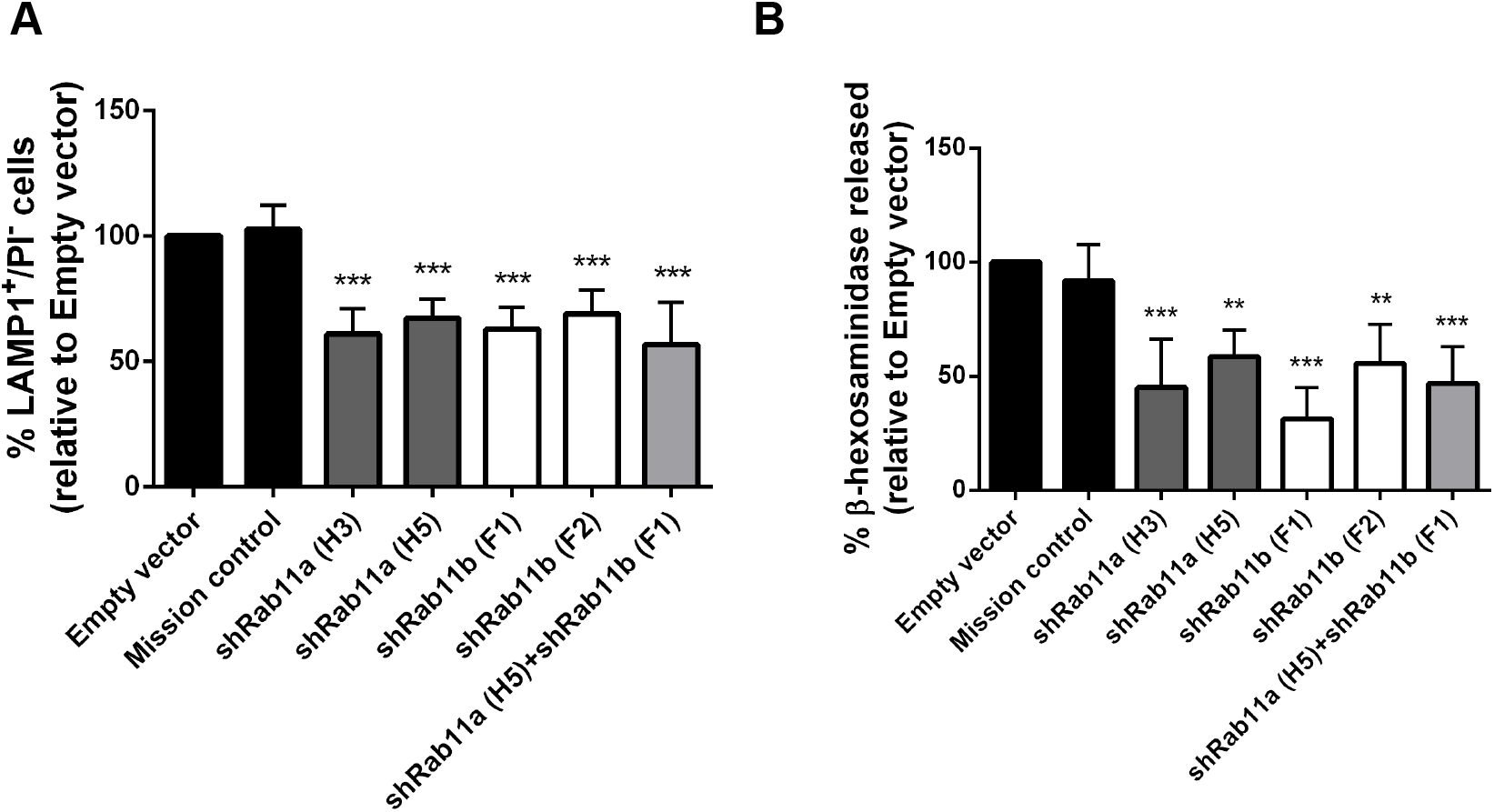
Rab11a or Rab11b silencing decreases LAMP1 cell surface expression and β-hexosaminidase release. HeLa cells transduced with lentiviruses encoding different shRNAs targeting Rab11a, Rab11b or both were treated with 10 μM ionomycin and 4 mM CaCl2 for 10 minutes at 37°C, to trigger lysosome exocytosis. (A) Cells were collected, stained with anti-LAMP1 antibody and analyzed by flow cytometry. Plot represents the percentage of LAMP1-positive cells and PI-negative cells. (B) Cell extracts and supernatants were collected and β-hexosaminidase release was quantified as described in Materials and Methods. Cells transduced with an empty vector (Empty) or a non-targeting shRNA (Mission) were used as negative controls. Results were normalized to the empty vector and are represe ted as mean ± SD of three independent experiments (***P<0.001, **P<0.01, *P<0.05).

To ascertain that Rab11a and Rab11b are specifically involved in lysosome exocytosis rather than in late endosome (LE) exocytosis, as LAMP1 is also present in these organelles, we analyzed the release of the lysosomal hydrolytic enzyme β-hexosaminidase, upon ionomycin stimulation. We observed that cells depleted for Rab11a or Rab11b display a significant impairment in the release of β-hexosaminidase, when compared with cells transduced with the empty vector or the vector encoding a non-targeting shRNA (Fig. 1B). Notably, HeLa cells transduced simultaneously with Rab11a and Rab11b shRNAs do not show a higher impairment in lysosome exocytosis, suggesting that the function of Rab11a and Rab11b in lysosome exocytosis is not redundant (Fig. 1A, B).

Next, we overexpressed GFP-tagged Rab11a or Rab11b and found that it does not lead to a significant change in LAMP1 cell surface expression levels and only slightly increases β-hexosaminidase release, upon ionomycin stimulation, when compared with cells transfected with the vector encoding only GFP (Fig. S2A, B). This result suggest that calcium-induced lysosome exocytosis cannot be further enhanced by overexpressing Rab11a or Rab11b, probably because there is a fixed lysosome pool that can undergo exocytosis. Surprisingly, HeLa cells overexpressing the constitutively-active mutants Rab11a Q70L/Rab11b Q70L or the dominant-negative mutants Rab11a S25N/Rab11b S25N do not show significant differences in LAMP1 cell surface expression levels or β-hexosaminidase release, when compared with cells overexpressing the wild-type form of the proteins (Fig. S2A, B). Thus, Rab11 GTP-binding state does not seem to affect calcium-dependent lysosome exocytosis in HeLa cells.

### Rab11-positive vesicles transiently interact with late endosomes/lysosomes at the cell tips, upon ionomycin stimulation

To gain insights into the molecular mechanism by which Rab11a and Rab11b regulate lysosome exocytosis, we analyzed the intracellular localization of Rab11a, Rab11b and LE/lysosomes, at steady-state and upon ionomycin stimulation, by immunofluorescence confocal microscopy. We observed that overexpressed GFP-Rab11a and GFP-Rab11b strikingly colocalize in HeLa cells (Fig. S3A). In addition, similar to the endogenous proteins, overexpressed GFP-Rab11a and GFP-Rab11b are distributed throughout the cytoplasm, with a striking accumulation at the perinuclear region, the typical localization of the endocytic recycling compartment (ERC) in these cells. Furthermore, the LE/lysosome marker LAMP1 accumulates in the perinuclear region but also localizes throughout the cytoplasm and at the cell tips, confirming our previous observations (Encarnação et al., 2016). The stimulation with ionomycin increases significantly the accumulation of LAMP1 at the cell tips, where it localizes in close proximity with Rab11a-positive vesicles (Fig. 2A). The same was observed for Rab11b (Fig. 2A). Thus, our results suggest that Rab11a- and Rab11b-positive vesicles interact with LEs/lysosomes at the cell periphery just before lysosome fusion with the plasma membrane. Furthermore, we observed that the silencing of Rab11a or Rab11b does not affect the distribution of LAMP1-positive vesicles in the cell (Fig. S3B).

**Fig. 2.**
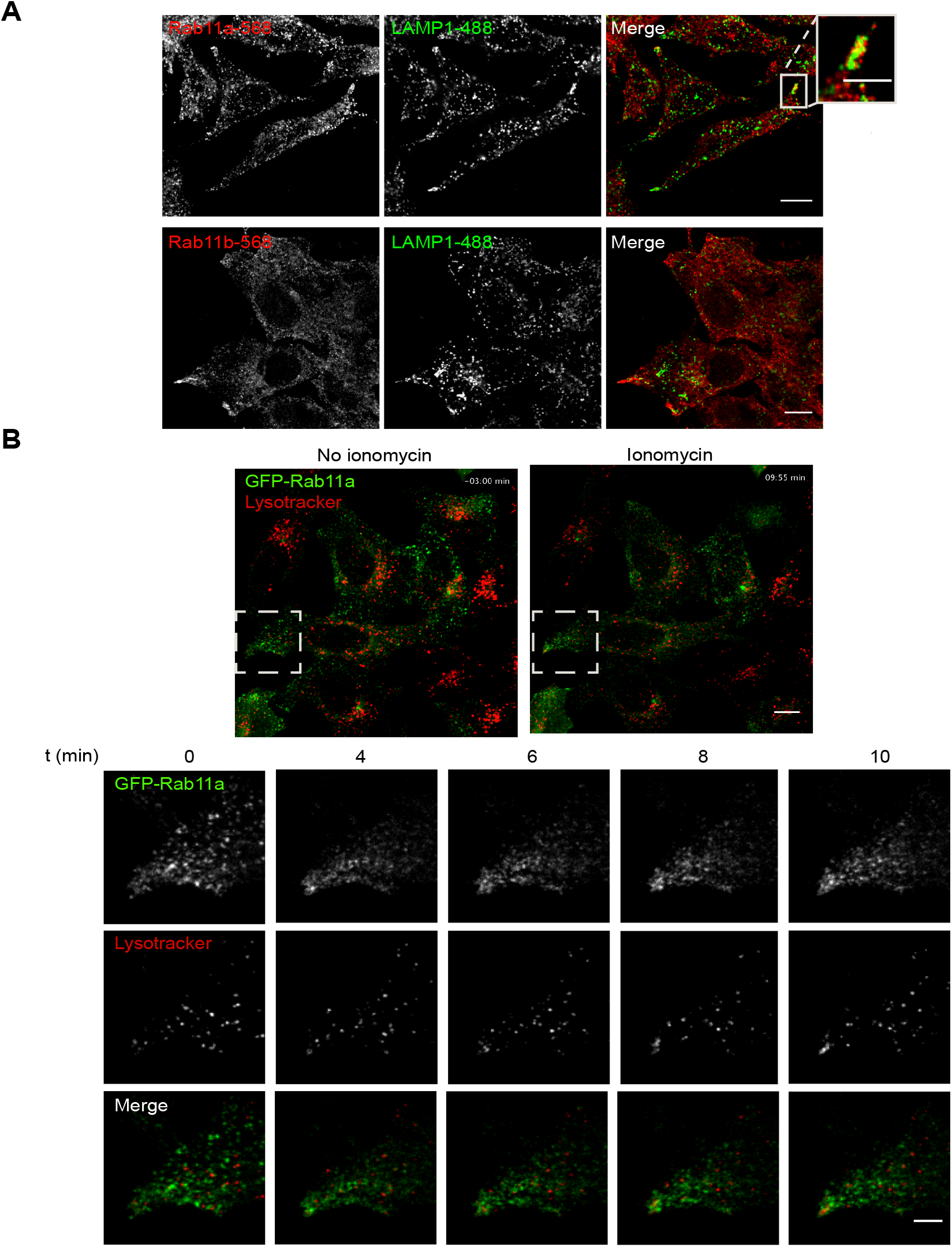
Rab11a transiently colocalizes with late endosomes/lysosomes at the cell tips, upon ionomycin stimulation. (A) Representative confocal microscopy images of the intracellular localization of Rab11a, Rab11b and LAMP1, in HeLa cells, upon lysosome exocytosis stimulation with 2.5 μM ionomycin and 4 mM CaCl2. Cells were fixed and stained with rabbit anti-Rab11a or anti-Rab11b (red) and mouse anti-LAMP1 (green) antibodies. White arrows indicate the areas at the cell tips where LAMP1 localize in close proximity with Rab11a- or Rab11b-positive vesicles. Scale bar: 10 μm. An enlarged view of a cell tip is shown in the inset. Scale bar: 5 μm. (B) Live cell imaging of HeLa cells transiently transfected with GFP-Rab11a (green) and incubated for 1-2 hours with LysoTracker (red) to labell lysosomes. The cells were imaged for 3 minutes before adding ionomycin and for 10 minutes after ionomycin stimulation. Images were captured every 5-10 seconds. Scale bar: 10 μm. The region outlined with a square was zoomed-in and the channels were split. Different time-frames: 0 min (immediately after adding ionomynin), 4, 6, 8 and 10 minutes are shown. Scale bar: 5 μm. Results represent two independent experiments.

It has been described that upon stimulation, only a small percentage of lysosomes undergo exocytosis (Jaiswal et al., 2002). Since this is a rare event, we used live cell imaging to detect transient interactions between Rab11-positive vesicles and lysosomes. For that, we transfected HeLa cells with GFP-Rab11a and tracked lysosomes labelled with LysoTracker, which stains acidic compartments. To understand the impact of ionomycin stimulation in Rab11a and lysosome intracellular localization, we imaged the cells for three minutes, at steady-state, and for ten minutes after ionomycin stimulation (Fig. 2B, Video 1). As observed in fixed cells, GFP-Rab11a accumulates at the perinuclear region, although it is also distributed throughout the cell (Fig. 2B, Video 1). Similarly, lysosomes labelled with LysoTracker localize near the perinuclear region, but are also found distributed throughout the cytoplasm. Upon ionomycin stimulation, the increase in intracellular calcium concentration leads to an accumulation of LysoTracker-positive vesicles at the cell tips, where they colocalize with GFP-Rab11a-positive vesicles for a few seconds before disappearing. Similar results were obtained upon overexpression of GFP-Rab11b (Video 2). Altogether, these results suggest that the increase in intracellular calcium concentration triggers the transport of lysosomes to the cell periphery, where they transiently interact with Rab11-positive vesicles before fusion with the plasma membrane.

### Sec15 is required for calcium-dependent lysosome exocytosis, independently of the exocyst tethering complex

To further understand the role of Rab11 in calcium-dependent lysosome exocytosis, we searched for Rab11 effector proteins that could mediate this function. To investigate if any of the known Rab11 effectors are involved, we silenced FIP1C and FIP2, two members of the Class I Rab11-FIP family, postulated to regulate the movement of recycling vesicles (Baetz and Goldenring, 2013; Schafer et al., 2014); Myosin Va and Myosin Vb, which are involved in vesicle transport along the actin cytoskeleton (Lapierre et al., 2001; Lindsay et al., 2013); and the exocyst subunits Sec15a and Sec15b, which are part of the octameric exocyst tethering complex (Wu et al., 2005; Zhang et al., 2004). Silencing efficiency was confirmed by qRT-PCR (Fig. S5A) and calcium-dependent lysosome exocytosis was evaluated by detecting LAMP1 at the cell surface, by flow cytometry (Fig. 3A), as well as measuring β-hexosaminidase release (Fig. 3B). Cells transfected with a non-targeting siRNA (siControl) were used as a control. Among the Rab11 effectors tested, we observed that only Sec15 reduces significantly LAMP1 cell surface expression levels, as well as β-hexosaminidase release, phenocopying the effect of Rab11a and Rab11b depletion. Curiously, LAMP1 cell surface expression levels are only decreased in the absence of Sec15b isoform but β-hexosaminidase release decreases when either Sec15a or Sec15b isoforms are silenced. Therefore, Sec15a isoform seems to play a more specific role in lysosome exocytosis, while Sec15b may regulate both LE and lysosome exocytosis. Noteworthy, the silencing of FIP1C, FIP2 or Myosin Vb has the opposite effect and leads to the increase of LAMP1 cell surface expression levels (Fig. 3A). This suggests that these proteins negatively regulate LE/lysosome exocytosis. The silencing of Myosin Vb also increases β-hexosaminidase release but, surprisingly, the silencing of Myosin Va decreases β-hexosaminidase release, suggesting that the two Myosin V isoforms play opposite roles in calcium-dependent lysosome exocytosis.

**Fig. 3.**
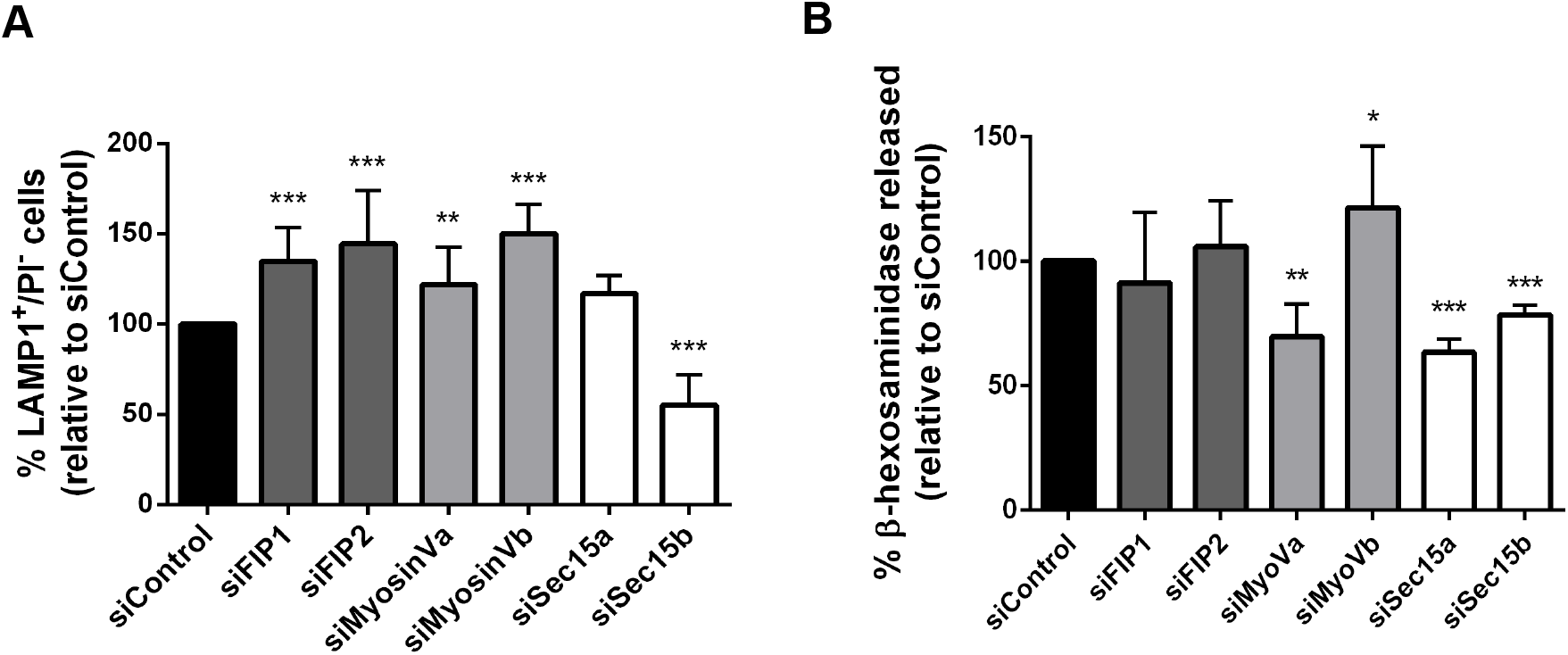
Rab11 effector proteins are involved in lysosome exocytosis. HeLa cells silenced for the indicated Rab11 effectors were treated with 10 μM ionomycin and 4 mM CaCl2 for 10 minutes at 37°C, to trigger lysosome exocytosis. (A) Cells were collected, stained with an anti-LAMP1 antibody and analyzed by flow cytometry. Plot represents the percentage of LAMP1-positive cells and PI-negative cells. (B) Extracts and supernatants of HeLa cells depleted of Rab11 effectors and treated with ionomycin were collected and β-hexosaminidase release was quantified as described in Materials and Methods. Cells transfected with a non-targeting siRNA (siControl) were used as a negative control. Results were normalized to the siControl and are represented as mean ± SD of three independent experiments (***P<0.001, **P<0.01, *P<0.05).

Since we wanted to find effector proteins that could mediate Rab11 function in lysosome exocytosis, we decided to further explore the role of Sec15, whose silencing has a similar effect to Rab11 depletion in lysosome exocytosis. Sec15 is part of the exocyst, a complex involved in the tethering of vesicles, including Rab11-positive recycling endosomes (Takahashi et al., 2012). To investigate if the exocyst complex is involved in the tethering and fusion of lysosomes to the plasma membrane upon stimulation, we silenced HeLa cells with siRNAs targeting different exocyst subunits. We analyzed Sec8, a core component of the exocyst; Sec10, which forms a subcomplex with Sec15, in yeast and mammalian cells (Guo et al., 1999; Moskalenko et al., 2003); and Exo70, which mediates the association of the exocyst to the plasma membrane (He et al., 2007). Silencing efficiency was confirmed by qRT-PCR (Fig. S1E). Surprisingly, the silencing of Sec8, Sec10 or Exo70 does not affect β-hexosaminidase release, when compared with a non-targeting siRNA (siControl) (Fig. 4A). Therefore, only the silencing of Sec15 subunit impairs lysosome exocytosis, independently of the exocyst complex. Noteworthy, we observed that GFP-Rab11 co-immunoprecipitates with myc-tagged Sec15a, in HeLa cells (Fig. 4B, Fig. S4A), as well with other exocyst subunits, namely Sec8 and Exo70 (Fig. S4B), thus confirming that Rab11 interacts with the exocyst tethering complex, as previously described by others (Takahashi et al., 2012). Next, we analyzed the intracellular localization of both Sec15a and Rab11. As expected, GFP-Sec15a extensively colocalizes with endogenous Rab11 (Fig. 4C). Moreover, Sec15a overexpression promotes endogenous Rab11 relocalization, increasing the number of Rab11-positive vesicles at the cell tips (Fig. 4C). This result is in agreement with previous reports, showing that Rab11 effectors, namely FIPs, influence Rab11 localization when overexpressed (Baetz and Goldenring, 2013). Overall, our results suggest that Sec15, together with Rab11, plays an important role in calcium-dependent lysosome exocytosis, and that the role of Sec15 is independent of the exocyst complex.

**Fig. 4.**
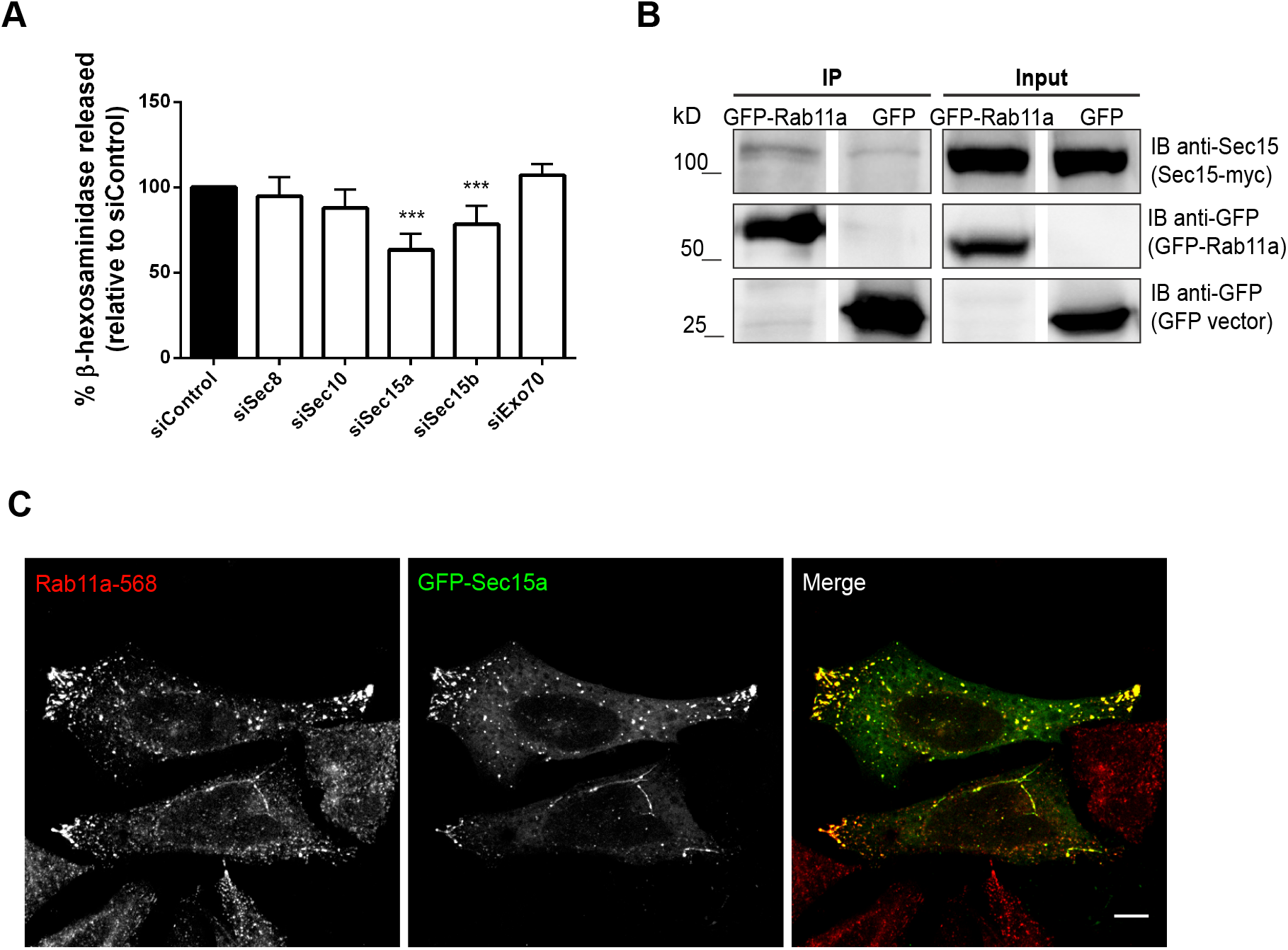
Sec15 is required for lysosome exocytosis independently of the exocyst complex. (A) HeLa cells silenced with siRNAs for the indicated exocyst subunits were treated with 10 μM ionomycin and 4 mM CaCl2 for 10 minutes at 37°C, to trigger lysosome exocytosis. Cell extracts and supernatants were collected and β-hexosaminidase release was quantified as described in Materials and Methods. Cells transfected with a non-targeting siRNA (siControl) were used as controls. Results were normalized to the siControl and are represented as mean ± SD of three independent experiments (***P<0.001, **P<0.01, *P<0.05). (B) Co-immunoprecipitation of GFP-Rab11a with Sec15a-myc in HeLa cells. Total cell extracts (500 μg) were used to immunoprecipitate Rab11a-GFP. Cells expressing GFP were used as a negative control. Input corresponds to 1/10 of total cell extracts used for IP (50 μg). Immunoblot was done using mouse anti--Sec15 and goat anti-GFP antibodies. Results represent two independent experiments. (C) Representative confocal microscopy images of HeLa cells stained with rabbit anti--Rab11a antibody (red) and overexpressing the exocyst subunit GFP-Sec15a (green). Scale bar: 10 μm. Results represent three independent experiments.

### Rab11-GRAB-Rab3a cascade is required for calcium-dependent lysosome exocytosis

Upon stimulation, Rab11-positive vesicles transiently interact with lysosomes near the cell periphery, immediately before lysosome fusion with the plasma membrane. Recently, we showed that Rab3a silencing impairs lysosome exocytosis (Encarnação et al., 2016). Rab3a was shown to recruit effector proteins, namely Slp4-a and NMIIA and regulate lysosome positioning and exocytosis. Moreover, a small percentage of lysosomes that localize close to the cell periphery were labeled with Rab3a. These likely correspond to the pool of lysosomes that fuses with the plasma membrane upon the increase in intracellular calcium concentration. Therefore, we hypothesized that Rab11 interacts with Rab3a on the surface of lysosomes. To test this hypothesis, we overexpressed GFP-Rab3a and mCherry-Rab11a, in HeLa cells, and performed co-immunoprecipitation studies. Upon immunoprecipitation of GFP-Rab3a, we detected a band corresponding to mCherry-Rab11a (Fig. 5A), indicating that these proteins do interact. Co-immunoprecipitation of mCherry-Rab11a using and anti-mCherry antibody followed by the detection of GFP-Rab3a (Fig. 5A) confirmed the interaction between these two Rab proteins.

**Fig. 5.**
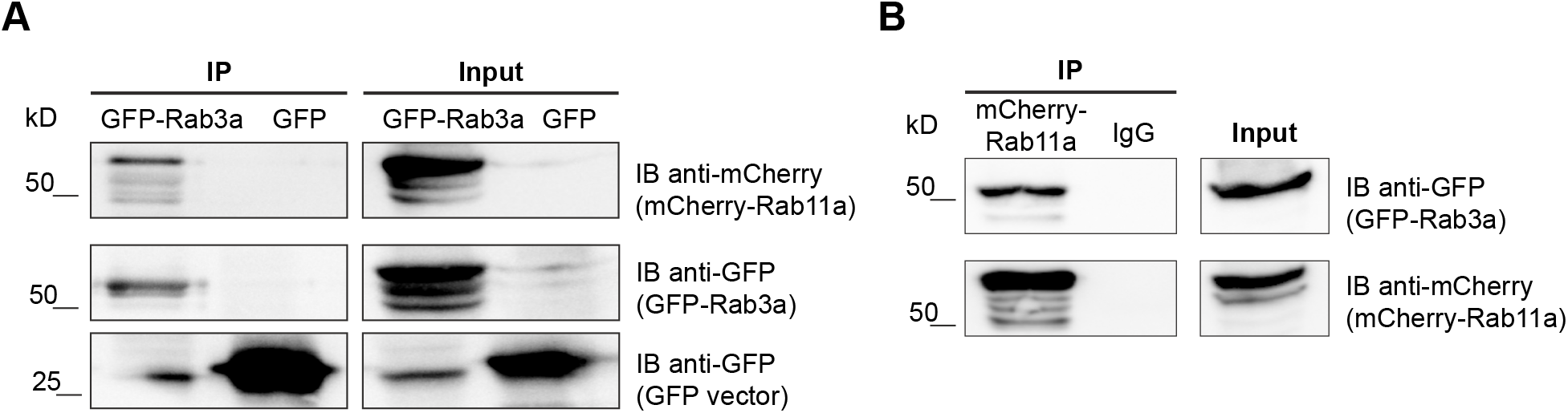
Rab11 co-immunoprecipitates with Rab3a. Total cell extracts (500-600 μg) were used to immunoprecipitate (A) GFP-Rab3a or (B) mCherry-Rab11a. GFP-transfected cells and goat IgG were used as negative controls. Immunoblot was done using goat anti-mCherry or goat anti-GFP antibodies to detect mCherry-Rab11a or GFP-Rab3a. Input corresponds to 1/10 of total cell extracts used for immnoprecipitation. Results represent three independent experiments.

Since there is no precedent for Rab proteins interacting directly, we postulated that Rab11 and Rab3a interaction is mediated by an adaptor protein. Searching for a protein that could link Rab3a and Rab11, we noticed that the Rab3a GEF GRAB (Luo et al., 2001) binds directly to Rab11a and Rab11b (Horgan et al., 2013). Since GRAB is a dual Rab binding partner, we investigated if GRAB could play a role in lysosome exocytosis through Rab11a and Rab3a. We started by confirming the interaction between Rab11a and GRAB, in HeLa cells. For that, we co-immunoprecipitated mCherry-Rab11a with wild-type (WT) GRAB-GFP or a mutant form of GRAB (GRAB∆223-GFP) with impaired Rab11 binding capacity (Horgan et al., 2013). As expected, WT GRAB co-immunoprecipitates efficiently with mCherry-Rab11a, whereas GRAB∆223-GFP shows weaker binding to Rab11a (Fig. 6A). Upon immunoprecipitation of WT GRAB-GFP or GRAB∆223-GFP it was also possible to detect Rab11a (Fig. S5A), confirming the interaction.

**Fig. 6.**
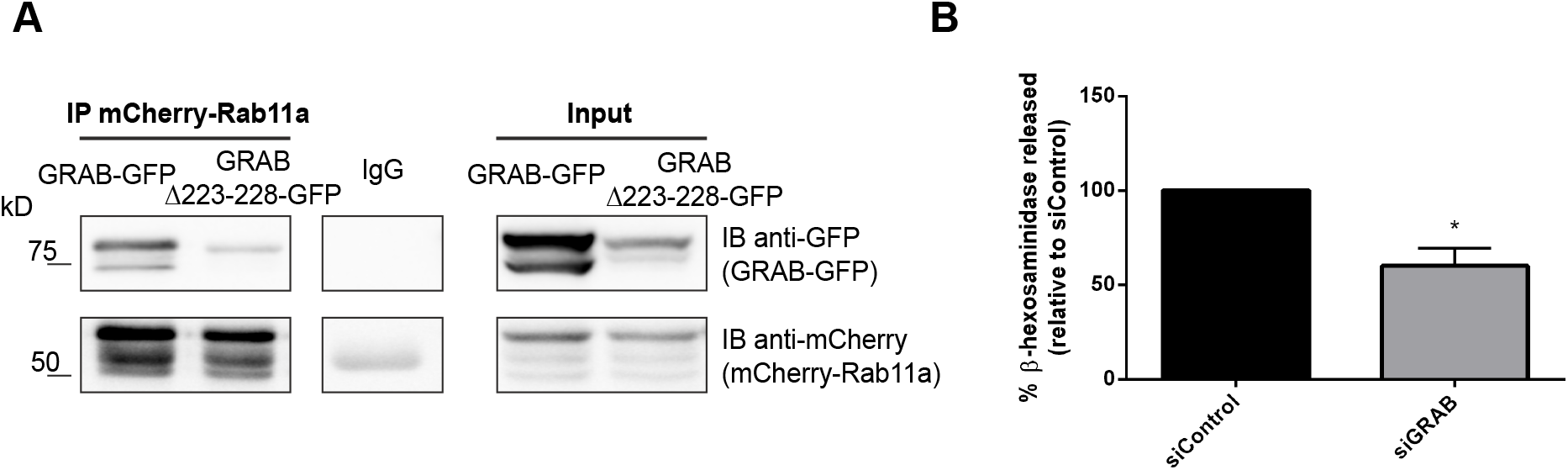
Rab11 interacts with Rab3a via GRAB. (A) Total cell extracts (600 μg) were used to immunoprecipitate mCherry-Rab11a using goat anti-mCherry antibody. Goat IgG was used as a negative control. Immunoblot was done using goat anti-GFP anti-body to detect WT GRAB-GFP or GRABΔ223-GFP. Input corresponds to 1/10 of total cell extracts used for immunoprecipitation. Results represent two independent experiments. (B) HeLa cells were transfected with siRNA for GRAB. β-hexosaminidase release was evaluated as described in Materials and Methods. Cells transfected with a non-targeting siRNA (siControl) were used as a control. Results were normalized to the siControl and are represented as mean ± SD of three independent experiments. (*P<0.05)

Next, we investigated if GRAB is required for lysosome exocytosis. To this end, GRAB was silenced using an siRNA pool and we monitored the release of β-hexosaminidase, upon ionomycin stimulation. Silencing efficiency was assessed by qRT-PCR (Fig. S5B). Interestingly, we found a significant decrease in β-hexosaminidase release when GRAB is depleted, as compared with a non-targeting siRNA (siControl) (Fig. 6B). This suggests that GRAB is also required for lysosome exocytosis, phenocopying Rab11 and Rab3a depletion. Furthermore, we were not able to clearly show an interaction between GFP-Rab3a or GRAB-GFP and myc-tagged Sec15a (Fig. S5C, D), suggesting that Rab11interaction with Sec15 could be lost before Rab11 interacts with GRAB and Rab3a.

## Discussion

Lysosome exocytosis is present in most cell types (Samie and Xu, 2014) and is required for several biological functions, including plasma membrane repair and lysosomal secretion (Appelqvist et al., 2013; Hämälistö and Jäättelä, 2016; Jaiswal et al., 2002; Reddy et al., 2001; Samie and Xu, 2014). Although several reports attempted to dissect the molecular players involved in this pathway, the role of Rab small GTPases is not well established. Recently, we screened the Rab family of small GTPases for their requirement in lysosome exocytosis and identified Rab3a and Rab10 as regulators of lysosome positioning and plasma membrane repair (Encarnação et al., 2016; Vieira, 2016). In the same screen, Rab11a and Rab11b, which have known roles in the regulation of endocytic recycling, were also identified as potentially involved in lysosome exocytosis. Curiously, Rab11b has been described by our group to be involved in the exocytosis of melanosomes, a type of LRO (Tarafder et al., 2014). More recently, Rab11a deficiency was shown to induce changes in LE/lysosomes, leading to an increased number of acidic vesicles labelled with LysoTracker localized to the perinuclear region (Zulkefli et al., 2019).

In this study, we confirmed and expanded the results obtained in the screen and show that Rab11a and Rab11b are key regulators of lysosome exocytosis, in HeLa cells. Indeed, the silencing of Rab11a or Rab11b isoforms significantly impairs lysosome exocytosis, in response to calcium stimulation. Interestingly, the reduction in lysosome exocytosis is not further enhanced when both Rab11 isoforms are silenced simultaneously, suggesting that both Rab11 isoforms are important but their function is not redundant.

To investigate the mechanism by which an endocytic recycling Rab protein could be involved in lysosome exocytosis, we used live cell imaging to track Rab11-positive vesicles and lysosomes. Upon ionomycin stimulation, we observed that Rab11-positive vesicles interact transiently with LE/lysosomes near the cell periphery. We postulate that this interaction is important for the delivery of molecules required for the last steps of lysosome tethering and fusion with the plasma membrane. Studies in cytotoxic T cells, have already shown the existence of transient interactions between immature lytic granules and Rab11-positive vesicles, suggesting them to be essential to prime lytic granules for the last steps of exocytosis (Ménager et al., 2007; Van der Sluijs et al., 2013). Therefore, in the absence of Rab11, lysosomes could lack some of the machinery required for efficient exocytosis, leading to lysosome secretion impairment.

Similar to other Rab GTPases, Rab11 regulates vesicular trafficking by recruiting effector proteins. Therefore, we searched for Rab11 effectors that could play a role in lysosome exocytosis. Although Rab11 has several known effector proteins (Baetz and Goldenring, 2013; Hales et al., 2001; Horgan and McCaffrey, 2009; Junutula et al., 2004; Lapierre et al., 2001; Lindsay et al., 2013; Wu et al., 2005; Zhang et al., 2004), we observed that from the ones tested, only the silencing of the exocyst subunit Sec15 leads to a consistent decrease in LAMP1 cell surface expression and β-hexosaminidase release, phenocopying Rab11 depletion. Additionally, confocal immunofluorescence microscopy confirmed that GFP-Sec15 colocalizes with endogenous Rab11 near the cell surface. This suggests that Sec15 acts together with Rab11 in the regulation of lysosome exocytosis. Sec15 is part of the exocyst complex, implicated in the tethering of secretory vesicles to the plasma membrane (Heider and Munson, 2013) and exocytosis of recycling endosomes (Takahashi et al., 2012). Recently, our group also demonstrated that the exocyst complex, together with Rab11b, is required for melanosome exocytosis (Moreiras et al., 2019). Surprisingly, in HeLa cells, the silencing of other exocyst subunits does not affect lysosome exocytosis, suggesting that the role of Sec15 in lysosome exocytosis is independent of the exocyst complex. Few studies have addressed the question of whether exocyst sub-complexes or exocyst subunits can perform functions independently of the exocyst complex (Moskalenko et al., 2003; Singh et al., 2019), but our results clearly suggest that exocyst subunits like Sec15, can perform functions independent of the complex.

Although the precise role of Sec15 in lysosome exocytosis is not entirely clear, it is likely that it could act as an adaptor protein between Rab11-positive vesicles and other molecules, for example Myosins. This would be important for the efficient trafficking of Rab11-positive vesicles to the cell periphery, where they interact with lysosomes and prime them for exocytosis. In fact, Sec15 has been shown to interact with the Myosin Va orthologue Myo2, in yeast (Jin et al., 2011) and *Candida albicans* (Guo et al., 2016). Furthermore, Sec15 was shown to participate in the transport of secretory vesicles by acting as an adaptor between Myo2 and the Rab11 orthologue Ypt31/32 or Rab8/10 homologue Sec4, in yeast (Jin et al., 2011). A similar mechanism could occur in mammalian cells. Although we were not able to show an interaction between Sec15 and Myosin Va, by co-immunoprecipitation (unpublished data), the silencing of Myosin Va also leads to a decrease in β-hexosaminidase release, suggesting a possible role for Myosin Va in lysosome secretion. Further studies are needed to elucidate if Sec15 interacts with MyosinVa in mammalian cells, and if a Rab11-Sec15-MysoinVa complex is required for lysosome exocytosis.

Since both Rab3a and Rab11 regulate lysosome exocytosis, we investigated if they interact. Indeed, by co-immunoprecipitation, we found that Rab11 interacts with Rab3a. Therefore, we speculate that Rab11-positive vesicles interact with the subset of Rab3a-positive lysosomes that localizes at the cell periphery and are prone to fuse with the plasma membrane. Although we cannot discard a direct interaction between Rab11 and Rab3a, Rab proteins usually interact via adaptor proteins. By searching the literature for molecules that could link the two Rab small GTPases, GRAB became an obvious candidate, since it is described to interact directly with Rab11 (Horgan et al., 2013) and it is also a Rab3a GEF (Luo et al., 2001). As hypothesized, we found that GRAB silencing leads to a decrease in β-hexosaminidase release and impairs lysosome exocytosis, phenocopying Rab11 and Rab3a depletion. Therefore, our data suggests that GRAB binds to Rab11 and is transported by Rab11-positive vesicles to the vicinity of Rab3a-positive lysosomes, where it acts as a Rab3a GEF, converting Rab3a to its active form (Fig. 7). Thus, we uncovered a Rab11-GRAB-Rab3a cascade essential for lysosome exocytosis. Interestingly, a similar cascade, involving Rab27-GRAB-Rab3a was recently described to regulate human sperm exocytosis (Quevedo et al., 2019). Furthermore, since we could not detect an obvious interaction between GRAB or Rab3a and Sec15, it is likely that Rab11 interacts with GRAB and Rab3a after interacting with Sec15, therefore suggesting that Sec15 plays a role in a different step of the lysosome exocytosis process (Fig. 7). Further studies should be done to elucidate this question.

**Fig. 7.**
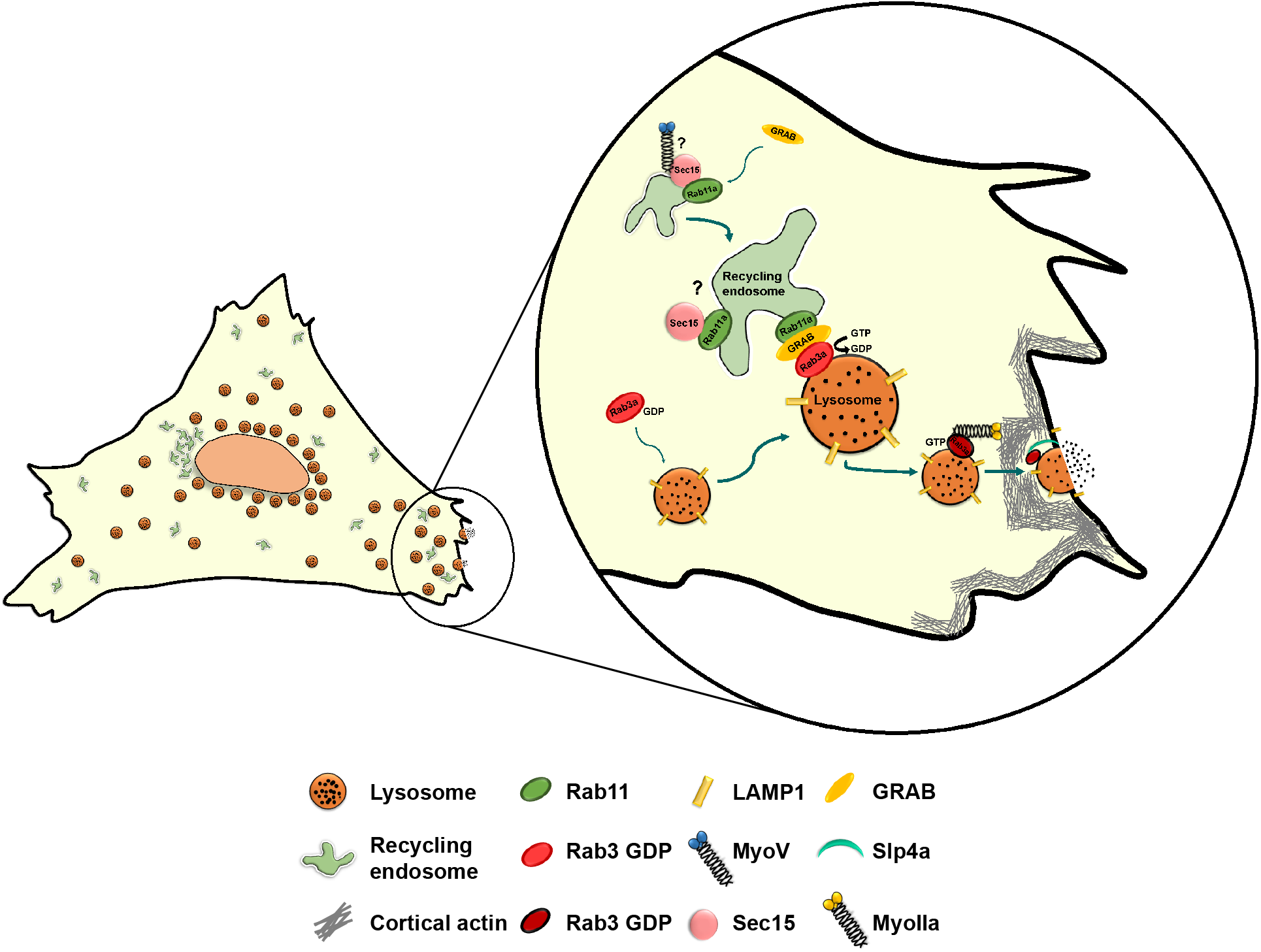
Proposed mechanism for the role of Rab11 in calcium-triggered lysosome exocytosis. Rab11 binds directly to Sec15, a subunit of the exocyst complex. The role of Sec15 role seems to be independent of the exocyst complex and could act as an adaptor protein between Rab11 and Myosins, facilitating the transport of Rab11-positive vesicles along the actin cytoskeleton. In this process, Rab11 possibly loses the interaction with Sec15 and interacts with GRAB, allowing the transport of Rab11-positive vesicles to the cell periphery. Once localized in the vicinity of the plasma membrane, Rab11-positive vesicles carrying GRAB interact with Rab3a-positive lysosomes. GRAB, can then activate Rab3a, which recruits NMIIA and Slp-4a, allowing the correct positioning lysosomes for lysosome exocytosis.

Thus, our study provides valuable insights into the role of the endocytic recycling pathway in calcium-triggered lysosome exocytosis. Moreover, the knowledge acquired can shed light on the molecular mechanism behind several human disorders that affect plasma membrane repair and LRO exocytosis.

## Materials and Methods

### Cell culture

HeLa and HEK293T cells were maintained at 37°C and 5% CO2, in Dulbecco’s modified Eagle’s medium (DMEM, Invitrogen) supplemented with 10% fetal bovine serum (FBS, Invitrogen), 100 U/ml penicillin G, 100 mg/ml streptomycin, 2 mM L-glutamine and 20 mM HEPES (Invitrogen). Both cells lines were obtained from the Division of Rheumatology, Immunology and Allergy, Brigham and Women’s Hospital, Harvard Medical School. These cell lines were not recently authenticated nor tested for contamination.

### Lentiviral production and transduction

Lentiviruses were produced in HEK293T cells in accordance with BL-2+ protocols publicly available at the Broad Institute’s RNAi Consortium Web site (www.broadinstitute.org/rnai/public/resources/protocols). VSV-G, pLKO.1, and pCMV-dR8.91 plasmids were obtained from the RNAi Consortium. The shRNA sequences are listed in Table S1. Controls used were Empty vector and Mission (CAACAAGATGAAGAGCACCAA) (Sigma).

For lentiviral transduction, HeLa cells plated in 6-well plates, were incubated with shRNA-encoding lentivirus supernatants in the presence of 6 μg/ml polybrene (hexadimethrine bromide; Sigma-Aldrich). Twenty-four hours after infection, virus-containing media were replaced by selection media containing 2 µg/ml puromycin (Sigma). All assays were performed 7–8 days after infection.

### Cell transfection

Gene silencing of Rab11 effector proteins was performed with siGENOME SMARTpool oligos (Thermo Scientific Dharmacon) using DharmaFECT 1 (Dharmacon), according to the manufacturer’s instructions. Briefly, HeLa cells, plated in 24 well-plates, were incubated with a transfection mixture containing 40 nM of siRNA and 2 µL of transfection reagent. Assays were performed 72-96 hours after siRNA transfection. The list of small interfering RNA (siRNA) sequences is described in Table S2. As a control, a non-targeting siRNA sequence, siControl (UAAGGCUAUGAAGAGAUAC) (Thermo Scientific Dharmacon) was used.

Plasmid transfections were performed using Lipofectamine 2000 (Invitrogen) in accordance with the manufacturer’s instructions. Briefly, HeLa cells, plated in 24-well plates, 10 cm dishes or Nunc™ Lab-Tek™ Chambered Coverglasses (ThermoFisher Scientific) were incubated with 1 μg DNA/2 μL Lipofectamine, 10 μg DNA/20 μL Lipofectamine or 0.5 μg/1 μL of Lipofectamine, respectively. Experiments were performed 24-48 hours after transfection. Plasmids used are described in Table S3.

### RNA extraction, cDNA production and real-time quantitative PCR

Total RNA was prepared with RNeasy Mini Kit (Qiagen), according to the manufacturer’s instructions. Reverse transcription into complementary DNA (cDNA) was achieved by incubating 1 μg of total RNA with 1 mM of dNTP mix (Thermo Scientific) and 0.3 μg/uL of random primers p(DN) (Roche) at 65°C for 5 min, followed by incubation with 2x first strand buffer (Invitrogen), 20 mM DTT (Invitrogen) and 40 U/μL of Recombinant Ribonuclease Inhibitor RNaseOUT (Invitrogen) at 25°C for 2 minutes. Finally, 50 U/μL of Superscript II Reverse Transcriptase (Invitrogen) were added and samples incubated at 25°C for 10 minutes, 42°C for 50 minutes and finally at 70°C for 15 minutes. qPCR was performed using the Fast Start Essential DNA Green Master (Roche) kit according to the manufacturer’s instructions. Analysis was done in a qPCR Roche Light Cycler. β-actin was used as an endogenous control to normalize the expression level of each gene analyzed. A complete list of primer sequences used for qPCR is displayed in Table S4.

### Calcium-induced lysosome exocytosis assay

Lysosome exocytosis was induced in HeLa cells using the calcium ionophore, ionomycin (Sigma-Aldrich), as previously described (Encarnação et al., 2016). Briefly, HeLa cells were incubated in Ca^2+^- and Mg^2+^-free ice-cold Hanks Balanced Salt Solution (HBSS, Gibco) with 2.5-10 μM ionomycin in the presence of 4 mM CaCl_2_, for 10 minutes at 37°C. Cells incubated with HBSS alone were used as a control. After incubation, cells were placed on ice and processed for flow cytometry, β-hexosaminidase release assay or confocal cell microscopy.

### Flow cytometry

Cells incubated in HBSS, with or without 10 μM ionomycin and 4 mM CaCl_2_, were placed on ice and detached with flow cytometry buffer (PBS, 1% FBS and 2 mM EDTA). Cells were stained with an antibody that recognizes the luminal epitope of the human LAMP1 protein (clone H4A3, BioLegend) conjugated with Alexa fluor 488 or 647 for 30 minutes, at 4°C. To exclude dead cells, samples were incubated with 0.5 μg/mL of propidium iodide (PI) (Sigma-Aldrich) just before acquisition. Acquisition was performed in a FACS Calibur or FACS CANTO II flow cytometer and at least 30,000 cells were analyzed using FlowJo version 10.1r7 software.

### β-hexosaminidase release assay

Cells incubated in HBSS, with or without 10 μM ionomycin and 4 mM of CaCl_2_, were placed on ice and cell supernatants containing released β-hexosaminidase were collected. In parallel, cells were lysed with 1% IGEPAL and further diluted 1:5 in dH_2_O. β-hexosaminidase activity was determined by incubating cell supernatants or lysates in a 96-well plate with 6 mM of the substrate 4-methylumbelliferyl- N-acetyl-β-d-glucosaminide (4-MU-β-D-GlcNAc; Glycosynth) resuspended in HBSS with 40 mM sodium citrate and 88 mM Na_2_PO_4_, pH 4.5, for 15 minutes at 37°C. Fluorescence was measured in a plate spectrofluorimeter Infinite F200 Pro reader (Tecan) at 365 nm excitation and 450 nm emission. The protein content from cell supernatants and lysates was determined, simultaneously, using the BCA protein assay kit (Pierce Laboratories), as described by the manufacturer. Absorbance was measured at 560 nm in the same plate reader. HBSS and 1% IGEPAL diluted 1:5 were used as controls. β-hexosaminidase (β-hex) activity was calculated for each sample normalizing to total protein amount as following: β-hex activity in supernatant = (fluorescence (365/450) - HBSS alone)/protein μg. β-hex activity in cell lysate = (fluorescence (365/450) - IGEPAL alone)/protein μg. Total β-hex activity = β-hex activity in supernatant + 5x β-hex activity in cell lysate. Finally, the percentage of β-hex release was calculated as following: β-hex release (% of total) = 100 × (β-hex activity in the supernatant/total β-hex activity).

### Confocal microscopy and live cell imaging

HeLa cells, grown on coverslips, were incubated in HBSS, with or without 2.5 μM ionomycin and 4 mM of CaCl_2_. Cells were fixed for 15 minutes in 4% paraformaldehyde (Alfa Aesar) and blocked/permeabilized for 30 minutes in PBS containing 1% BSA and 0.05% saponin. Cells were then incubated with primary antibodies, namely rabbit anti-Rab11a (Abcam) (1:500), rabbit anti-Rab11b (Abgent) (1:100), mouse anti-LAMP1 conjugated with Alexa fluor 488 (BioLegend) (1:500) for 1 h, followed by incubation with Alexa-conjugated secondary antibodies goat anti-rabbit 488 or 568 (1:1000) (Invitrogen). Cells were further incubated with 1 μg/mL DAPI (Sigma) and mounted in Mowiol mounting medium (Calbiochem). Images were acquired on a Zeiss LSM 710 confocal microscope with a Plan-Apochromat 63×1.4 NA oil-immersion objective. Digital images were analyzed with LSM Image software or ImageJ.

For live-cell confocal imaging, HeLa cells were seeded in Nunc™ Lab-Tek™ Chambered Coverglasses (ThermoFisher Scientific) and transfected with GFP-Rab11a, as previously described. Twenty-four hours after transfection, cells were incubated with 50 nM LysoTracker^®^ Red DND-99 (Invitrogen) for 1-2 hours. After extensive washing with PBS, cells were kept in phenol red-free DMEM (Gibco) for imaging. Cells were imaged before and after stimulation with 4 μM ionomycin and 4 mM CaCl_2_. Live-cell imaging was performed in HBSS at 37°C using an Andor Revolution spinning disk confocal microscope (Andor Technology) equipped with an EMC CD camera with a Plan Apo VC PFS 60x objective (1.4 NA oil-immersion; Nikon). The system was controlled by iQ software (Andor Technology) and images were analyzed using ImageJ software.

### Immunoprecipitation

HeLa cells were lysed in ice-cold RIPA lysis buffer (50 mM Tris-HCl pH 7.5; 1 mM EDTA; 1 mM EGTA; 150 mM NaCl; 2 mM MgCl_2_; 1 mM DTT; 1% IGEPAL), in the presence of protease and phosphatase inhibitors, for 30 minutes, at 4°C. After centrifugation at 18,800 × *g* for 30 minutes, at 4°C, protein concentration was determined using the DC protein assay kit (Bio-Rad), according to the manufacturer’s instructions.

Proteins fused with GFP were immunoprecipitated for 2h at 4°C, using GFP-Trap Beads (ChromoTek), previously equilibrated in 150 mM NaCl RIPA, using 350-800 μg of total cell extracts. Samples were centrifuged at 2,500 × *g* for 3 minutes at 4°C and washed twice with RIPA containing 500 mM NaCl, and three times with RIPA containing 150 mM NaCl. Finally, samples were solubilized in 2x Laemmli sample buffer and boiled at 95°C for 5 minutes.

Immunoprecipitation of endogenous and mCherry-fused proteins was performed using 500-700 μg of total protein pre-cleared for 1 h with Protein G-Sepharose beads (GE Healhtcare Life Sciences). Cell lysates were incubated overnight, at 4°C, with 2 μg of goat anti-mCherry (Sicgen) or rabbit anti-Rab11a (Abcam) antibodies. The same concentration of goat IgG (#026202, Invitrogen) or rabbit IgG_1_ (I5006, Sigma) were used as negative controls. ProteinG-Sepharose beads were then added and incubated for 5 hours at 4°C, under constant agitation. Beads were recovered by centrifugation, washed twice in RIPA containing 500 mM NaCl, and three times in RIPA with 150 mM NaCl. Finally, samples were solubilized in 2x Laemmli sample buffer and boiled at 95°C for 5 minutes.

### Immunoblotting

Proteins were loaded on 8-10% SDS-polyacrylamide gels, transferred to Nitrocellulose membranes (GEHealtcare Life Sciences) for 60 minutes at 100V and processed for immunoblotting. Membranes were blocked with blocking buffer (5% milk in PBS with 0,1% Tween-20) and incubated with antibodies diluted in the same buffer. Primary antibodies used are described in Table S5. Horseradish peroxidase (HRP)-conjugated secondary antibodies (GE Healthcare) used were anti-mouse/rabbit (1:5,000) or anti-goat antibodies (1: 10,000) (Amersham ECL Goat, Rabbit or Mouse IgG, HRP-linked whole Ab, GE Healtcare Life Sciences). HRP activity was detected with Amersham ECL Select (GE Healtcare Life Sciences), according to the manufacturer’s instructions. Chemiluminescence was detected using a ChemiDoc™ Touch Imaging and the results were analyzed using ImageLab software.

### Statistical analysis

Numerical data are presented as mean ± standard deviation (SD). Unpaired, two-tailed Student’s t-test was used to compare different data sets with a control. Statistical analysis was performed using GraphPad Prism version 6.05.

## Abbreviations used in this paper

FIP: Rab11-family of interacting protein
GEF: guanine nucleotide exchange factor
LE: late endosomes
LRO: lysosome-related organelle
NMIIA: non-muscle myosin heavy chain IIA
Slp-4a: synaptotagmin-like protein 4a

## Acknowledgments

We would like to thank the cell culture, flow cytometry and microscopy facilities of CEDOC; Wei Guo and Mary MacCaffrey for the kind gift of plasmids; and Hugo Moreiras and Cristina Casalou for assistance in the preparation of Fig. 7. This study was supported by Fundação para a Ciência e Tecnologia (FCT): CE was supported by a post-doctoral fellowship (SFRH/BPD/78491/2011), LBL by a PhD fellowship (SFRH/BD/131938/2017) and DCB by the FCT Investigator Program (IF/00501/2014/CP1252/CT0001).

## Notes

### Competing Interest Statement

The authors have declared no competing interest.

