## Supplementary data for "A novel Rab11-Rab3a cascade required for lysosome exocytosis"

**Supplementary information:**

**Table S1** – shRNAs used to transduce HeLa cells.

| Gene name | Accession number | Hairpin number | Source/Clone ID | Target sequence (5′–3′) |
|---|---|---|---|---|
| ***RAB11A*** | NM_004663 | H2 | TRCN0000073018 | CCC TAT AAA CAT AGC AGG ATT |
| ***RAB11A*** | NM_004663 | H3 | TRCN0000073019 | CGA GCT ATA ACA TCA GCA TAT |
| ***RAB11A*** | NM_004663 | H4 | TRCN0000073020 | CCT GTC TCG ATT TAC TCG AAA |
| ***RAB11A*** | NM_004663 | H5 | TRCN0000073021 | GAG CTA TAA CAT CAG CAT ATT |
| ***RAB11A*** | NM_004663 | H6 | TRCN0000073022 | GCC TTA TTG GTT TAT GAC ATT |
| ***RAB11B*** | NM_004218 | F1 | TRCN0000029184 | GCC TTG GAT TCC ACT AAC GTA |
| ***RAB11B*** | NM_004218 | F2 | TRCN0000029185 | CCT ATT CAA AGT GGT GCT CAT |
| ***RAB11B*** | NM_004218 | F3 | TRCN0000029186 | CCA CTA ACG TAG AGG AAG CAT |
| ***RAB11B*** | NM_004218 | F4 | TRCN0000029187 | CGT AGA GGA AGC ATT CAA GAA |
| **RAB11B** | NM_004218 | F5 | TRCN0000029188 | CAA GCA CCT GAC CTA TGA GAA |
| **RAB11B** | NM_004218 | F5 | TRCN0000029188 | CAA GCA CCT GAC CTA TGA GAA |

**Table S2** – siRNAs used to silence Rab11 effectors.

| Gene name | Accession number | Target sequences |
|---|---|---|
| ***RAB11 FIP1*** | NM_001002814 | CAAACAGAAGGAAACGAUA |
| | | GCUAACUGCAGCUUGGGAA |
| | | AGUGAGAACUUGAACAAUG |
| | | UCCGCGAGCUGGAAGACUA |
| ***RAB11 FIP2*** | NM_001330167 | GGAUGAAGGUGAAUUGUGU |
| | | CGAGCUACCUGGAUUGCUA |
| | | GGAACAAUAUGACCGCAAG |
| | | GAUAAGAUGAAGGGUAGAA |
| ***MYOSIN VA (MYO5A)*** | NM_000259 | GAACAAAUGUGCACUCUUU |
| | | GAUCAUCUGCUCUGGAUUA |
| | | AAAGUAAGGUCGUUGCUAA |
| | | CGCAGGAGGUACAAGAUUA |
| ***MYOSIN VB (MYO5B)*** | NM_001080467 | GGACUUACCUCUUGGAGAA |
| | | CAAGUUGGCCUAACAGUGA |
| | | GCAGAUCUGGCCUACAAUA |
| | | ACAGUGGCCUUUAUACGAA |
| ***SEC8 (EXOC4)*** | NM_021807 | GAAUUGAGCAUAAGCAUGU |
| | | UAACUGAGUACUUGGAUAU |

| | | |
|---|---|---|
| | | GCCGAGUUGUGCAGCGUAA<br>ACUGAGUGACCUUCGACUA |
| ***SEC10 (EXOC5)*** | NM_006544 | GAAGUCCGAUGCAGAGCAA<br>GGAGAUACCUUAUGACACA<br>GGAAAGAAUUAGACAGCGU<br>CAUUAGGAGUGGAUCGGAA |
| ***SEC15A (EXOC6)*** | NM_019053 | GAAGUUUGGUGAAUGGUAU<br>GUUGAUGGCUAUAGAAGAU<br>GAUAGAGACAGUCGUGAAA<br>CCAAACUCCGUGAGGAUAU |
| ***SEC15B (EXOC6B)*** | NM_015189 | UAACUGAACUGCUGAAAGU<br>GUACUAGUCCGAAGUCUGA<br>CAAGUAAGCCACUAUCGAU<br>CGGGAAACAUUUGAGAAUU |
| ***EXO70 (EXOC7)*** | NM_001013839 | GGUUAAAGGUGACUGAUUA<br>GACCUUCGACUCCCUGAUA<br>CUAAGCACCUAUAUCUGUA<br>CGGAGAAGUACAUCAAGUA |
| ***GRAB (RAB3IL1)*** | NM_013401 | GAGAGAAGGGCUCCGAGUU<br>GGCCUGGACUUCACGGCAA<br>AGAUCAUGAGGUUGCGGAA<br>ACAUGAAGCAGGCGGCAUC |

**Table S3** – DNA plasmids used for transfection.

| Protein | Plasmid | Source |
|---|---|---|
| **GFP** | pENTR-GFPC1 | José Ramalho (Sicgen) |
| **mRab11a** | pENTR-GFPC1 | José Ramalho (Sicgen) |
| **mRab11b** | pENTR-GFPC1 | José Ramalho (Sicgen) |
| **mRab11a S25N** | pENTR-GFPC1 | José Ramalho (Sicgen) |
| **mRab11a Q70L** | pENTR-GFPC1 | José Ramalho (Sicgen) |
| **mRab11b S25N** | pENTR-GFPC1 | José Ramalho (Sicgen) |
| **mRab11b Q70L** | pENTR-GFPC1 | José Ramalho (Sicgen) |
| **mCherry** | pCDNA-ENTR-BP-mCherry | José Ramalho (Sicgen) |
| **mRab11a** | pCDNA-ENTR-BP-mCherry | José Ramalho (Sicgen) |
| **mRab11b** | pCDNA-ENTR-BP-mCherry | José Ramalho (Sicgen) |
| **mRab3a** | pENTR-GFPC2 | José Ramalho (Sicgen) |
| **rSec15** | pJ3 Myc-EGFP | Kind gift of Wei Guo |
| **rSec15** | pCDNA3 myc | Kind gift of Wei Guo |
| **GRAB** | pGFP | Kind gift of Mary McCaffrey |
| **GRAB∆223-228** | pGFP | Kind gift of Mary McCaffrey |

**Table S4** – Primers used for qRT-PCR.

| Target gene | Sequence |
|---|---|
| *β-ACTIN* | Forward 5'-GCAAAGACCTGTACGCCAAC-3' |
| | Reverse 5'-AGTACTTGCGCTCAGGAGGA-3' |
| *RAB11A* | Forward 5'-CGATGGCTGAAAGAACTGAG-3' |
| | Reverse 5'-GACAGCACTGCACCTTTGGC-3' |
| *RAB11B* | Forward 5'-TCACCCGCAACGAGTTCAAC-3' |
| | Reverse 5'-CTGCACCACGGTAGTACGC-3' |
| *FIP1* | Forward 5'-CAAGGAGCGAGGAGAAATTG-3' |
| | Reverse 5'-GGTGTCTGACCCACTGTCCT-3' |
| *FIP2* | Forward 5'-TGGGGGATCTGATAGCCCTT-3' |
| | Reverse 5'-ACTCATATGAAAACTTGAAGATGGC-3' |
| *SEC8* | Forward 5'-ATGGCCAGCAAGCACTATCT-3' |
| | Reverse 5'-AGGTGCCGGTGTAGTTCATC-3' |
| *SEC10* | Forward 5'-ATAAAGCAGTGCCAGGAGGG3' |
| | Reverse 5'-AGCCAGGACTGTTTCTGGATT3' |
| *SEC15A* | Forward 5'-GTCACTACACCGGAGCTCAA3' |
| | Reverse 5'-TGCTCCAGGTGTGTTGTGTT3' |
| *SEC15B* | Forward 5'-CTGACCCTGCTTGAGAAGATGA3' |
| | Reverse 5'-GCCACGGTGTCAATGAGTTTC-3' |
| *EXO70* | Forward 5'-CATGGGTTATCAGGGGATTTG-3' |

| | |
|---|---|
| | Reverse 5’-GAGGTCCAGGTGTGGGTAGA-3’ |
| *MYOSIN VA* | Forward 5’-CAGTGGTCAGAACATGGGTG-3’ |
| | Reverse 5’-TCGCATGGCATACTTAGCTG-3’ |
| *MYOSIN VB* | Forward 5’-AGAACTGGAGGAGGAGCGAT-3’ |
| | Reverse 5’-GGTTTGATGGGTTCCGCCTA-3’ |
| *GRAB* | Forward 5’-ATCCGCTACATCCAGCAAGG3’ |
| | Reverse 5’-TAAGCCTCCTGGGGGAAGAA3’ |

**Table S5** – Antibodies used for immunoprecipitation and immunoblotting.

| Primary antibodies | Host animal | Immunoblotting dilution |
|---|---|---|
| **Rab11a (Abcam) (ab128913)** | Rabbit | 1:50,000 |
| **Rab11b (Abgent) (AP12943b)** | Rabbit | 1:1,000 |
| **Sec8 (Enzo Life Sciences) (ADI-VAM-SV016-F)** | Mouse | 1:500 |
| **Sec15 (Sigma) (SAB4200612)** | Mouse | 1:1,000 |
| **Exo70 (Milipore) (MABT186)** | Mouse | 1:1,000 |
| **GFP (Sicgen) (AB0020-200)** | Goat | 1:1,000 |
| **mCherry (Sicgen) (AB0040-200)** | Goat | 1:1,000 |
| **GFP (NeuroMab) (clone N86/8)** | Mouse | 1:3,000 |

FigureS1

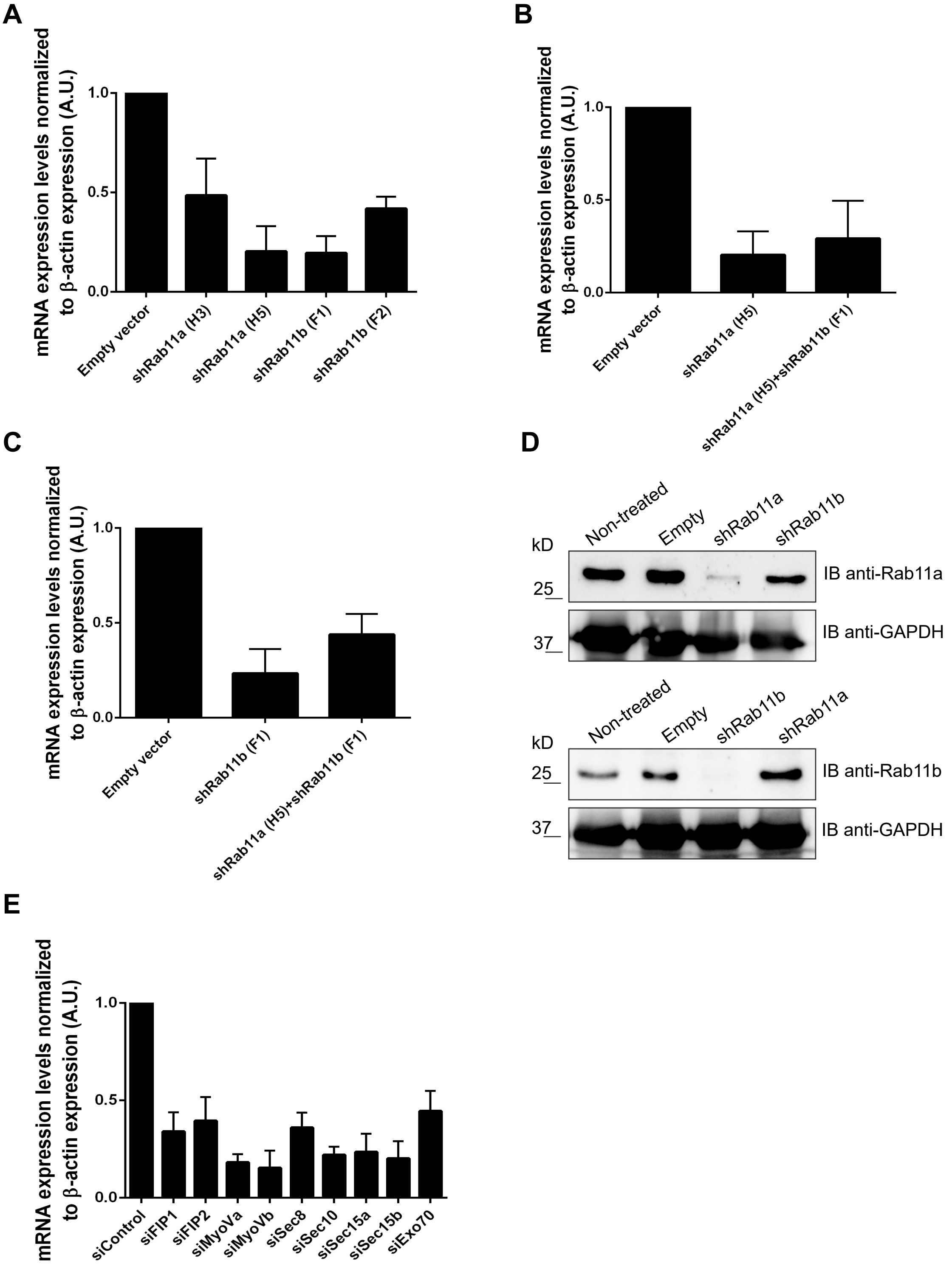


**Fig. S1. Rab11a, Rab11b and Rab11 effectors silencing efficiency.** (A) HeLa cells transduced with lentiviruses encoding different shRNAs targeting Rab11a (H3, H5) or Rab11b (F1, F2) were selected for 7 days with puromycin and silencing efficiency was evaluated by qRT-PCR. Results were normalized to the expression of β-actin (B) Rab11a expression was analyzed by qRT-PCR, as described before, when silenced alone or in combination with Rab11b. (C) Rab11b expression was analyzed by qRT-PCR, as described before, when silenced individually or in combination with Rab11a. All results were normalized to the empty vector and are represented as arbitrary units (A.U). Plots represent the mean ± SD of three or more independent experiments. (D) Rab11a and Rab11b protein expression levels were analyzed in HeLa cells transduced with lentiviruses encoding shRNA for Rab11a or Rab11b. Immunoblot was done using rabbit anti-Rab11a or anti-Rab11b antibodies. Non-transduced cells or cells transduced with empty vector were used as controls. Protein levels were normalized to GAPDH. The images are representative of two or more independent experiments. (E) Rab11 effectors were silenced in HeLa cells as described in Materials and Methods. The relative expression of each silenced gene was analyzed by qRT-PCR and normalized to the expression of β-actin. All results were normalized to siControl and are represented as arbitrary units (A.U). Results are representative of three independent experiments.

FigureS2

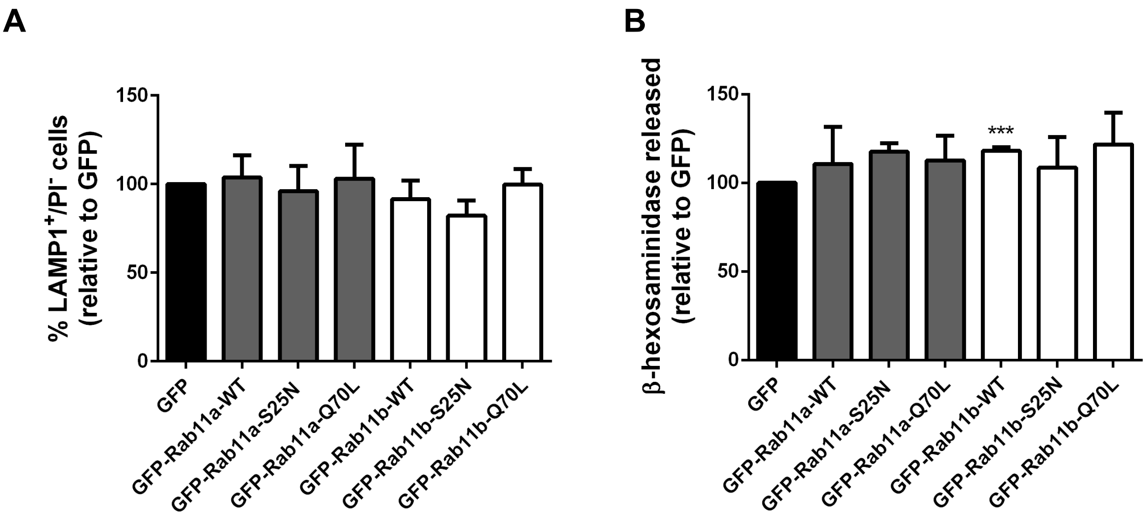


**Fig. S2. Overexpression of Rab11a or Rab11b wild-type or mutant forms does not affect LAMP1 cell surface expression or β-hexosaminidase release.** (A) HeLa cells overexpressing the wild-type (WT), dominant negative (S25N) or constitutively active (Q70L) mutants of GFP-Rab11a or -Rab11b were treated with 10 µM ionomycin and 4 mM CaCl2 for 10 minutes at 37ºC, to trigger lysosome exocytosis. Cells were collected, stained with an anti-LAMP1 antibody and analyzed by flow cytometry. Plots represents the percentage of LAMP1-positive cells and PI-negative cells. (B) Cellular extracts and supernatants were collected and β-hexosaminidase release was quantified as described in Materials and Methods. Results were normalized to the cells transfected with a vector encoding GFP alone and are represented as mean ± SD of three independent experiments.

FigureS3

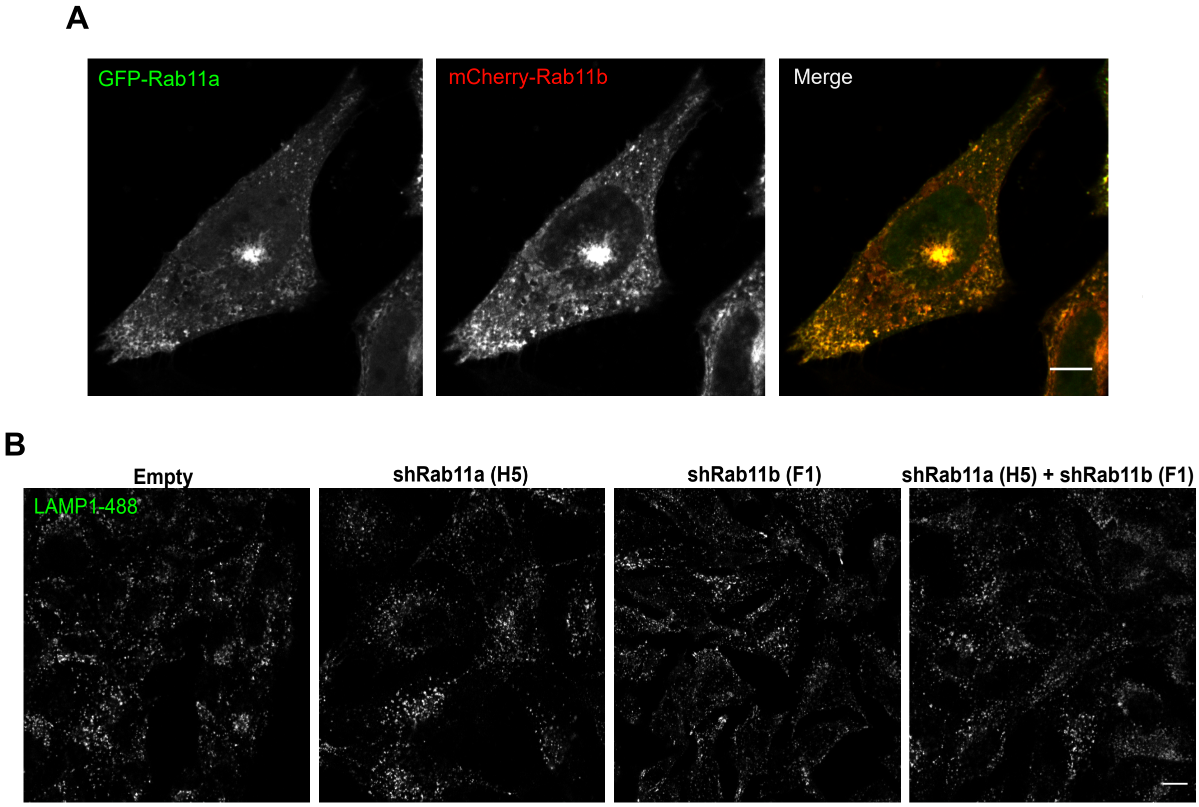


**Fig. S3. Rab11a and Rab11b display similar intracellular localizations and LAMP1 distribution is not affected upon Rab11a or Rab11b silencing.** (A) Representative confocal microscopy images of the localization of Rab11a and Rab11b in HeLa cells transfected with both GFP-Rab11a (green) and mCherry-Rab11b (red) plasmids. Scale bar: 10 μm. (B) Confocal microscopy images of the intracellular localization of LAMP1, in HeLa cells transduced with an empty vector (Empty) or lentiviruses encoding shRNAs targeting Rab11a (H5), Rab11b (F1) or both. Scale bar: 20 μm. Results are representative of three independent experiments.

FigureS4

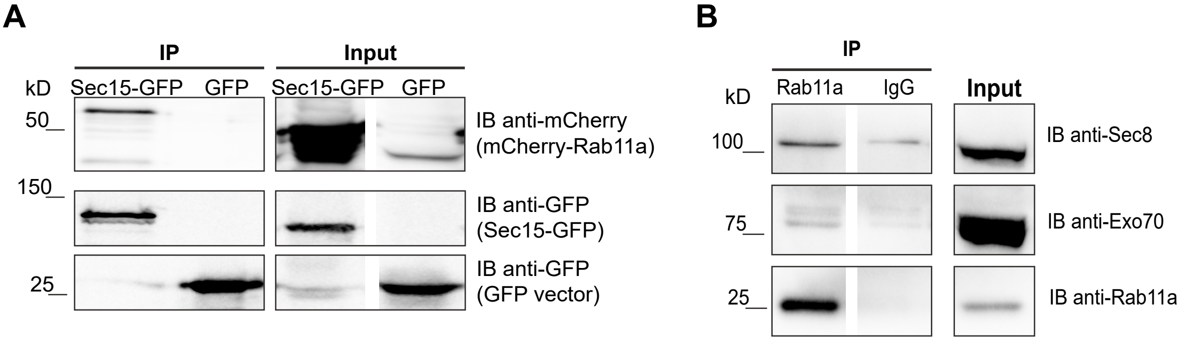


**Fig. S4. Rab11a co-immunoprecipitates with different exocyst complex subunits in HeLa cells.** (A) Total cell extracts (800 μg) were used to immunoprecipitate Sec15-GFP. Cells expressing GFP were used as a negative control. Immunoblot was done using goat anti--mCherry antibody to detect mCherry-Rab11a or goat anti-GFP antibody. (B) Total cell extracts (700 μg) were used to immunoprecipitate Rab11a, using rabbit anti-Rab11a antibody that recognizes specifically this isoform. Rabbit IgG was used as a negative control. Immunoblot was done using mouse anti-Sec8, mouse anti-Exo70 or rabbit anti-Rab11a antibodies. The images are representative of two or more independent experiments. Inputs correspond to 1/10 of total cell extracts used for immunoprecipitation. Results are representative of three independent experiments.

FigureS5

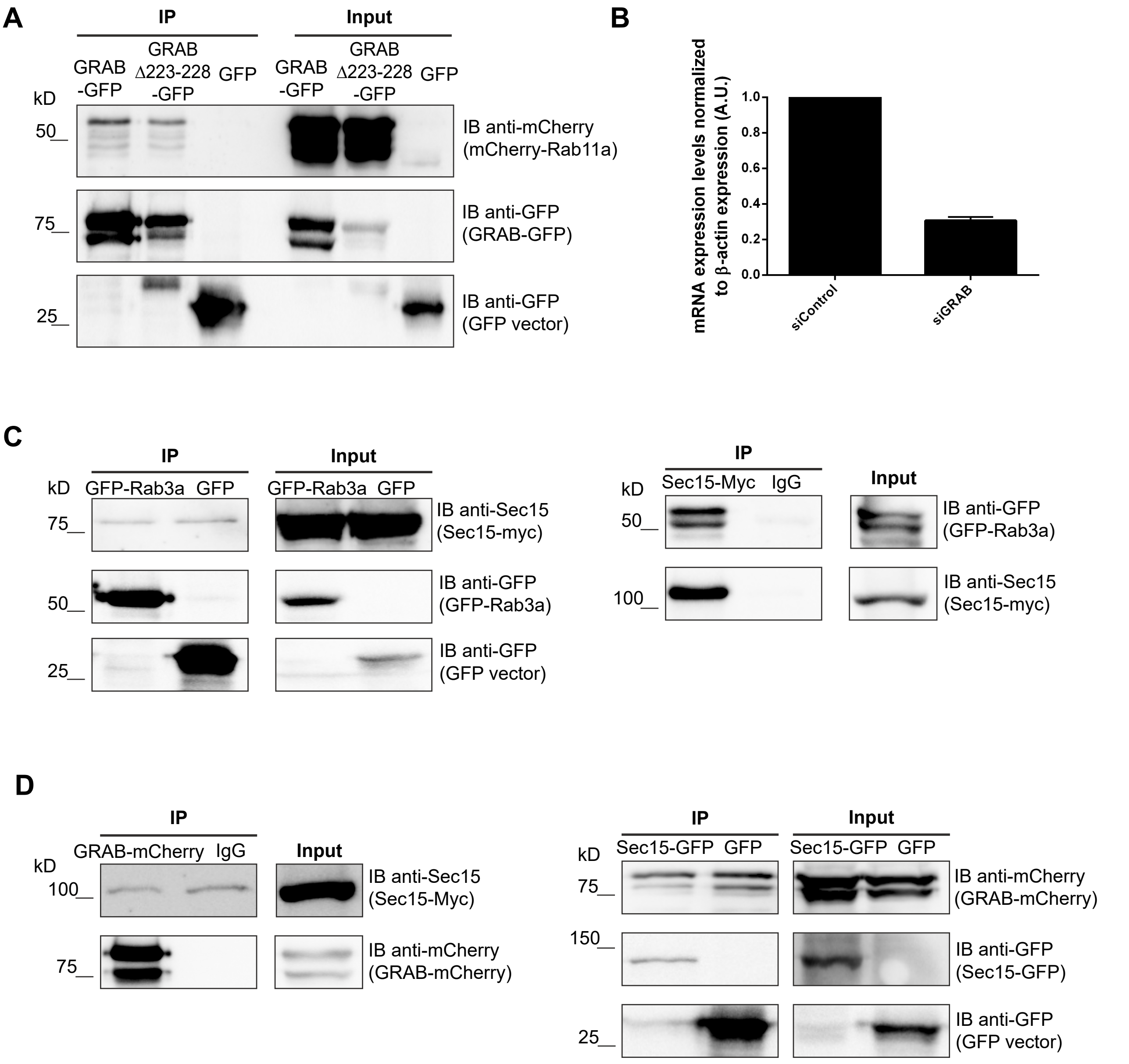


**Fig. S5. GRAB is required for Rab11a-Rab3a interaction.** (A) Total cell extracts (350 µg) were used to immunoprecipitate WT GRAB-GFP or GRABΔ223-GFP. GFP-transfected cells were used as a negative control. Immunoblot was done using goat anti-mCherry antibody to detect mCherry-Rab11a. Input corresponds to 1/10 of total cell extracts used for immunoprecipitation. (B) GRAB was silenced in HeLa cells as described in Materials and Methods. Relative mRNA expression was analyzed by qRT-PCR and normalized to the expression of β-actin. Results are represented as arbitrary units (A.U). (C) Total cell extracts (350 µg) were used to immunoprecipitate GFP-Rab3a or Sec15-Myc, respectively. GFP-transfected cells or Mouse IgG were used as negative controls. Immunoblot was done using goat anti-GFP or mouse anti-Sec15 antibodies to detect GFP-Rab3a or Sec15-Myc, respectively. Input corresponds to 1/10 of total cell extracts used for immunoprecipitation. (D) Total cell extracts (350-450 µg) were used to immunoprecipitate GRAB-mCherry or Sec15-GFP, respectively. GFP-transfected cells or Goat IgG were used as negative controls. Immunoblot was done using goat anti-GFP, goat anti-mCherry or mouse anti-Sec15 antibodies to detect Sec15-GFP, GRAB-mCherry or Sec15-Myc, respectively. Input corresponds to 1/10 of total cell extracts used for immunoprecipitation . Results are representative of three independent experiments.

**Video 1.** Live cell imaging of HeLa cells transiently transfected with GFP-Rab11a (green) and incubated for 1-2 hours with LysoTracker (red) to label lysosomes. The cells were imaged for 3 minutes before adding ionomycin (Time 0) and for 10 minutes after ionomycin stimulation. Images were captured every 5-10 seconds. The videos are representative of two independent experiments.

**Video 2.** Live cell imaging of HeLa cells transiently transfected with GFP-Rab11b (green) and incubated for 1-2 hours with LysoTracker (red) to label lysosomes. The cells were imaged for 3 minutes before adding ionomycin (Time 0) and for 10 minutes after ionomycin stimulation. Images were captured every 5-10 seconds. The videos are representative of two independent experiments.
